# Cerebellum metastasis model of HER2-positive breast cancer unveils key role of IL34-induced Arg1+ macrophages

**DOI:** 10.1101/2025.06.17.660224

**Authors:** Xiaoqing Cheng, Khooshbu Kantibhai Patel, Brandon Zhou, Yiwei Fu, Ryan Cleary, Maureen Highkin, Jacob Hsia, Xiaohua Jin, Benjamin Kohn, Julie L. Prior, Megan S. Michie, Rui Sun, Amanda E. D. Van Swearingen, James D. Quirk, Ian S. Hagemann, Tristan Li, Vaibhav Jain, Simon Gregory, Albert Kim, Ron Bose

**Affiliations:** Division of Oncology, Department of Medicine, Washington University School of Medicine, St. Louis, MO 63110; Duke Molecular Physiology Institute, Duke University School of Medicine, Durham, NC, 27710, USA; Department of Neurological Surgery, Washington University School of Medicine, St. Louis, MO 63110; Mallinckrodt Institute of Radiology, Washington University School of Medicine, St. Louis, MO 63110; Duke Center for Brain and Spine Metastasis, Duke University School of Medicine, Durham, NC, 27710, USA; Duke Cancer Institute, Duke University School of Medicine, Durham, NC, 27710, USA; Alvin J. Siteman Cancer Center, Washington University School of Medicine, St. Louis, MO 63110; Department of Pathology & Immunology, Washington University School of Medicine, St. Louis, MO 63110; Department of Neuroscience, Washington University School of Medicine, St. Louis, MO 63110; Hope Center for Neurological Disorders, Washington University School of Medicine, St. Louis, MO 63110; Department of Genetics, Washington University School of Medicine, St. Louis, MO 63110; The Preston Robert Tisch Brain Tumor Center, Duke University School of Medicine, Durham, NC, 27710, USA; Department of Neurosurgery, Duke University School of Medicine, Durham, NC, 27710, USA; The Brain Tumor Center, all at Washington University School of Medicine, St. Louis, MO 63110

**Author notes:** **Corresponding authors:** Ron Bose, Washington University School of Medicine, Division of Oncology, Mail Stop 8076-0041-03. 660 S. Euclid Ave. St. Louis, MO 63110.

**Keywords:** Breast cancer, brain metastasis, cerebellum, macrophage, HER2 positive breast cancer, spatial transcriptomics, Arg1, Interleukin-34, inflammation, trastuzumab

## Abstract

Brain metastases occur in up to 40% of Stage IV breast cancer patients. The cerebellum is a frequent location for metastases in HER2-positive breast cancer patients, but the mechanisms for this are unknown. Here, we developed a syngeneic, immunocompetent mouse model for breast cancer brain metastases by stereotactically injecting mouse HER2-overexpressing breast cancer organoids into the cerebellum. Growth of these cerebellar metastases was monitored by MRI and trastuzumab optical imaging using a near-infrared fluorophore conjugated to trastuzumab. Spatial transcriptomics identified interleukin-34 production by breast cancer cells inducing ARG1+ macrophages at the invading edge of the metastasis. Treatment with a blocking antibody to interleukin-34’s receptor, CSF1R, produced tumor shrinkage. These findings have immediate translation potential as a CSF1R-blocking antibody is FDA-approved. Further, it demonstrates that cancer-associated inflammation bordering the brain metastasis promotes metastatic growth and offers a molecularly targeted strategy to treat inflammation in brain metastasis.

**Graphical summary:** 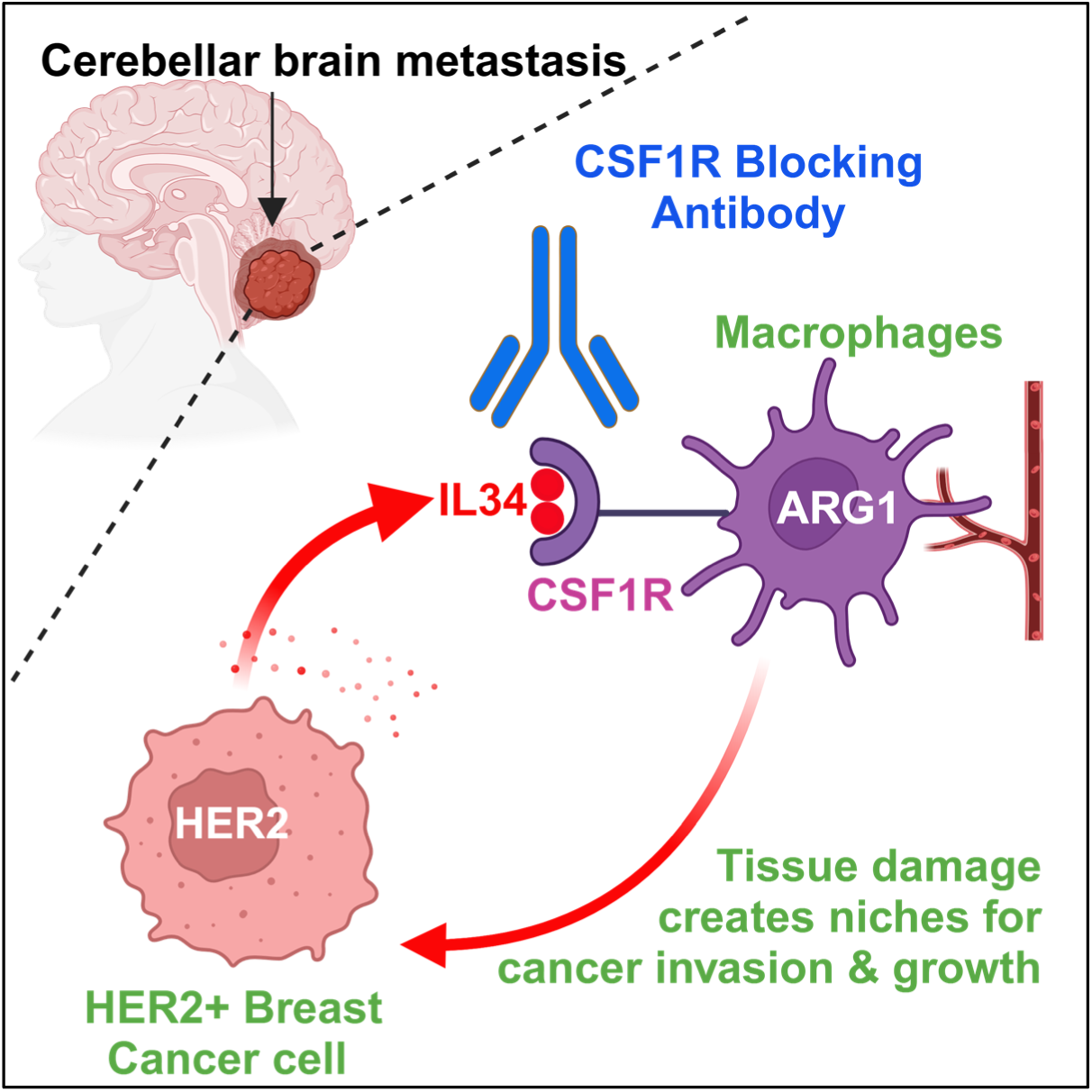

## INTRODUCTION

Breast cancer brain metastases (BCBM) are an emergency and possibly catastrophic event for breast cancer patients. Up to 40% of stage IV breast cancer patients will develop BCBM.^1, 2^ Two aggressive subtypes of breast cancer, HER2+, and triple-negative breast cancer, are the most likely to cause BCBM, but they do so with different clinical patterns.^1^ Triple-negative BCBM tend to be distributed throughout the brain,^3^ and they tend to occur when the patient also has cancer progression throughout the body, both inside and outside the CNS. In contrast, HER2+ BCBM has an increased predilection to metastasize to the cerebellum and will often occur when breast cancer outside of the CNS is under good control.^3, 4^ HER2+ BCBM can metastasize to other brain regions, but strikingly, in the German BCBM registry, 60% of the patients with HER2+ BCBM had metastases to the cerebellum.^4^ Cerebellum metastases pose high risk to patients because of their proximity to the fourth ventricle and brainstem, which can cause obstructive hydrocephalus or affect breathing and other essential body functions. The differences in the presence or absence of metastatic growth outside the CNS are attributed to the effect of trastuzumab and other HER2-targeted monoclonal antibodies, which are highly effective outside the CNS but cannot readily penetrate the intact blood-brain-barrier (BBB).^5^ However, the mechanisms causing the differential localization to different brain regions between HER2+ and triple-negative BCBM are unknown.

To model cerebellar BCBM from HER2+ breast cancer, we developed an immunocompetent, syngeneic transplantation strategy where breast cancer organoids obtained from a C57BL/6J (B6) mouse are injected into another B6 mouse. B6 mice are widely used for immunology research, and numerous transgenic and knock-out strains are available in this strain’s background. While FVB strain mice have traditionally been favored in breast cancer research, the differences in breast cancer development between B6 and FVB strains are small.^6^ Transplanting organoids between syngeneic mice provides an opportunity to separately genetically manipulate the “seed and soil” ^7^ of cerebellar BCBM. The “seed” is the breast cancer cell or organoid in this case, and the “soil” is the microenvironment of the cerebellar BCBM. While liver metastasis has been studied using transplanted human colorectal cancer organoids,^8^ to our knowledge, this is the first study to transplant genetically engineered murine organoids to study BCBM.

The microenvironment of the cerebellum has many features in common with other regions of the brain, but it also has unique elements specific to the cerebellum. Glial cell populations include astrocytes, oligodendrocytes, and microglia, but in the cerebellum, transcriptionally defined subpopulations of these cell types specific to the cerebellum have been described.^9^ The neuronal cell types of the cerebellum are unique, with Purkinje neurons notable for their large dendritic trees and small granule cell neurons, which are highly numerous. In fact, the cerebellum constitutes only 10% of the brain mass, but contains more than 50% of the brain’s neurons, due to the very large number of these small, granule cell neurons.^10^ In addition to these resident cell types of the cerebellum, neuroinflammation or other stimuli can recruit bone marrow derived macrophages to the cerebellum. During BCBM initiation and growth, changes in many of these cell types have been described. Reactive glia are seen in and around BCBM, with reactive astrocytes showing hypertrophy and fewer dendritic branches compared to their normal morphology.^11^ Astrocytes stimulated with Type I interferon can recruit monocytic myeloid cells to the BCBM.^12^ A major hypothesis in the field is that breast cancer cells alter the brain microenvironment and that this altered brain-tumor microenvironment contributes to further growth of the BCBM.^11^ Understanding the interactions occurring in the microenvironment of the cerebellum when HER2+ BCBM develop and grow is the major focus of this study.

In this study, we developed a model for HER2+ BCBM to the cerebellum in an immunocompetent mouse and performed spatial transcriptomics to characterize the brain-tumor microenvironment. We identified interleukin-34 (IL34) secretion by the breast cancer cells inducing ARG1+ macrophages to the periphery of the metastasis where tumor invasion occurs. Because CSF1R is the receptor for IL34, we treated mice with established cerebellar BCBM with a CSF1R blocking antibody and observed shrinkage of the brain metastases. These findings have immediate translational potential because, in August 2024, the FDA approved a blocking CSF1R monoclonal antibody, axatilimab, for the treatment of another cancer-related condition, chronic graft versus host disease.^13^ Our findings suggest that this CSF1R antibody could potentially also benefit brain metastases patients.

## RESULTS

### Organoid transplant model for HER2-positive BCBM to the cerebellum

Figure 1A shows a patient with two large HER2+ BCBM in the cerebellum. Remarkably, this patient did not have any metastases elsewhere in the brain. Her case and that of other similar patients motivated us to develop an animal model of cerebellar HER2+ BCBM to better understand, treat, and hopefully someday prevent this type of metastasis. An accurate preclinical model of HER2+ breast cancer cerebellum metastases is urgently needed to help us understand the molecular mechanisms of brain metastases and the tumor cell-intrinsic factors within the cerebellar microenvironment.

**Figure 1.**
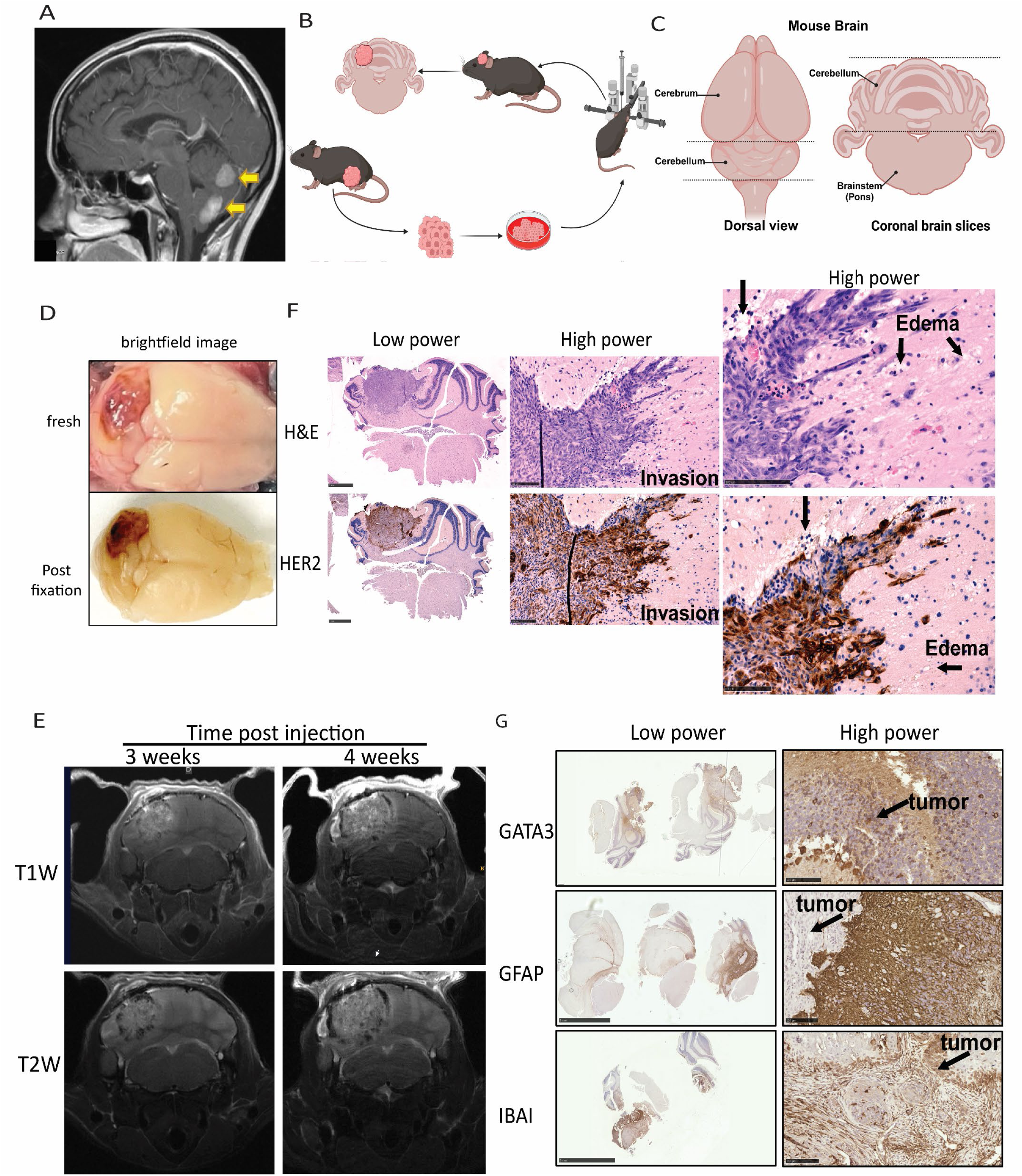
Organoids transplant model for HER2 positive breast cancer cereblleum metastases. (A) MRI of HER2+ patient shows specific cerebellum metastasis in the brain. (B) Overview of work flow to establish breast cancer brain metastasis model. (C) Dorsal and coronal brain slices view of mouse cerebellum. (D) Brightfield images of dissected and post-fixed cerebellum tumor. (E) T1-weighted and T2-weighted MRI of stereotaxic injection at 3 weeks, 4 weeks post-transplantation of HER2+ organoids in C57BL/6 syngeneic mice. (F) Top: H&E staining distinguishes dense tumor growth characteristics of primary epithelial tumors from normal brain parenchyma. Along the border of the lesion, areas of invasion and edema with microvacuolization can be seen. Bottom: IHC stained with ErbB2 antibodies confirm the overexpression of HER2 in the intracranial lesion, as well as HER2 expression in the invading cells. Scale bars of lower power images are 1 mm. Scale bars of high-power images are 100 µm. (G) IHC stained with GATA3, GFAP, and IBA1 antibodies confirm the properties of breast cancer and distinct features of cerebellum tumors. Scale bars of lower power images are 5 mm. Scale bars of high-power images are 500 µm.

An intracranial injection strategy (Figure 1B) was chosen because of its feasibility and anatomical precision. Intracranial injection is widely used to study glioblastoma multiforme (GBM) and in studies with BCBM using human patient derived xenografts (PDX).^14^ Using stereotactic injection equipment, the location and brain region of the BCBM can be precisely controlled. Further, this direct injection approach yields very predictable timing and a high success rate for metastasis development. While intracardiac injection models have been favored for studying BCBM,^11, 15–17^ it is more challenging for studying cerebellar BCBM for several reasons. The blood supply to the cerebellum predominantly comes from the vertebral artery and the posterior circulation of the brain.^10^ Intracardiac injection delivers most of the breast cancer cells to the cerebrum and not to the cerebellum. Similarly, carotid artery injection primarily does not reach the posterior circulation of the brain. Developing a breast cancer cell line that homed specifically to the cerebellum would be desirable, but is not currently available.^18^ The ideal model would be a transgenic mouse that spontaneously develops cerebellar BCBM, but unfortunately, no such mouse model currently exists, to our knowledge.

We previously developed a transgenic mouse that overexpresses the human HER2 gene with a V777L activating mutation.^19, 20^ The use of HER2 transgenes with activating mutations is very common in the HER2 field. The first HER2 transgenic mouse made carries the rat HER2 V664E activating mutation driven by the MMTV promoter.^21^ More recently, a Tet-inducible HER2 transgenic mouse also used this same rat HER2 V664E activating mutation.^22^

We chose the human HER2 gene for our transgenic construct because trastuzumab is species specific, recognizing human, but not rodent HER2.^19, 20^ Further, our human HER2^V777L^ transgene is driven by an actin promoter, generating over-expression with intense circumferential membrane staining that is clinically scored as 3+ IHC staining.^20, 23^

Our transgenic mouse has three additional important properties. First, it contains a Lox-STOP-Lox element generating conditional expression.^19^ Second, we found that the addition of a second genetic alteration, either *PIK3CA*^H1047R^ activating mutation or *TP53* deletion, dramatically accelerates tumorigenesis, matching genotypes seen in human cancer patients.^20^ Third, we found that breast cancer organoids derived from these mice can be re-implanted into the mammary fat pad of syngeneic B6 mice and will form mammary tumors that spontaneously metastasize to the lungs of the recipient mice.^20^ Therefore, we hypothesized that injection of these organoids into other sites in syngeneic mice could create models of other types of breast cancer metastases.

Figure 1B shows the schema used to generate cerebellar BCBM in mice. We used B6 mice expressing HER2^V777L^; *PIK3CA*^H1047R^ (HP) activating mutations or HER2^V777L^; *TP53^flox/floxl^* (H53) In brief, breast cancer primary tumors were established in either HP or H53 mice by injections of Cre-expressing adenovirus into the mammary gland, these tumors were harvested, and murine breast cancer organoid cultures were established from these tumors. These murine cancer organoids were luciferase labeled and expanded *in vitro.* For the intracranial injections, HP or H53 organoids were dissociated to small clusters, and they were stereotactically injected unilaterally into the cerebellum (Figure 1B). One thousand or five thousand cells were injected per mouse. Figure 1C shows the location of the cerebellum on the dorsal view of the intact mouse brain and in coronal brain slices or images. Weeks later, after confirming implantation and BCBM growth by MRI, the mice were sacrificed. After dissection and fixation, the cerebellum tumor had a hemorrhagic appearance distinct from the normal brain tissue (Figure 1D).

We used MRI imaging to monitor the tumor metastatic status *in vivo* (Figure 1E). Axial T1-weighted post-contrast MRI of the cerebellum revealed a hyperintense lesion with irregular borders at the site of injection that grew in the cerebellum over 4 weeks (Figure 1E). At 3 weeks post-injection, the lesion was located above the 4th ventricle and brainstem. While gradual growth was seen from 3 weeks post-injection, the tumor mass was significantly larger at 4 weeks post-injection, exerting pressure on the 4th ventricle (Figure 1E).

Following histological sectioning and staining, we identified several areas of interest around the lesion (Figure 1F). Areas of invasion could be identified at the border of the BCBM. Tumor cells there were depolarized and had a flattened cell morphology, resembling a motile phenotype (Figure 1F, middle panels). In addition, areas of edema and microvacuolization were present at the edge of the metastasis (Figure 1F, black arrows), indicative of cancer-associated inflammation bordering the metastasis. To validate that the primary tumor subtype had been retained, the tumor was stained with HER2 antibody, and HER2 over-expression was confirmed (Figure 1F, bottom). Most importantly, cells in areas of invasion retained their HER2 expression. Overall, this animal model provides a feasible approach to study cerebellar BCBM and we characterized it further.

### Immunohistochemical Characterization of HER2+ Cerebellar BCBM

We performed IHC staining on these BCBM for several markers. GATA3 is a transcription factor expressed by breast cancer cells and is present in the BCBM (Figure 1G). GFAP is a marker of astrocytes and other glial cell types, including ependymal cells. GFAP expression is absent in these BCBMs (Figures 1G, 2A-2B). IBA1 (Ionized calcium-binding adaptor molecule 1), also called allograft inflammatory factor 1 (AIF1), is a well-established marker for microglia/macrophages. IBA1-positive cells were seen within the BCBM (Figure 1G). The HP and H53 organoids are estrogen receptor α (ERα) positive^20^ (Figure S1), and ERα expression is seen in these BCBM (Figure 2A-B). About half of human HER2+ BCBM are ER+,^2^ therefore these HER2+ ER+ BCBM’s recapitulate what occurs in patients.

**Figure 2.**
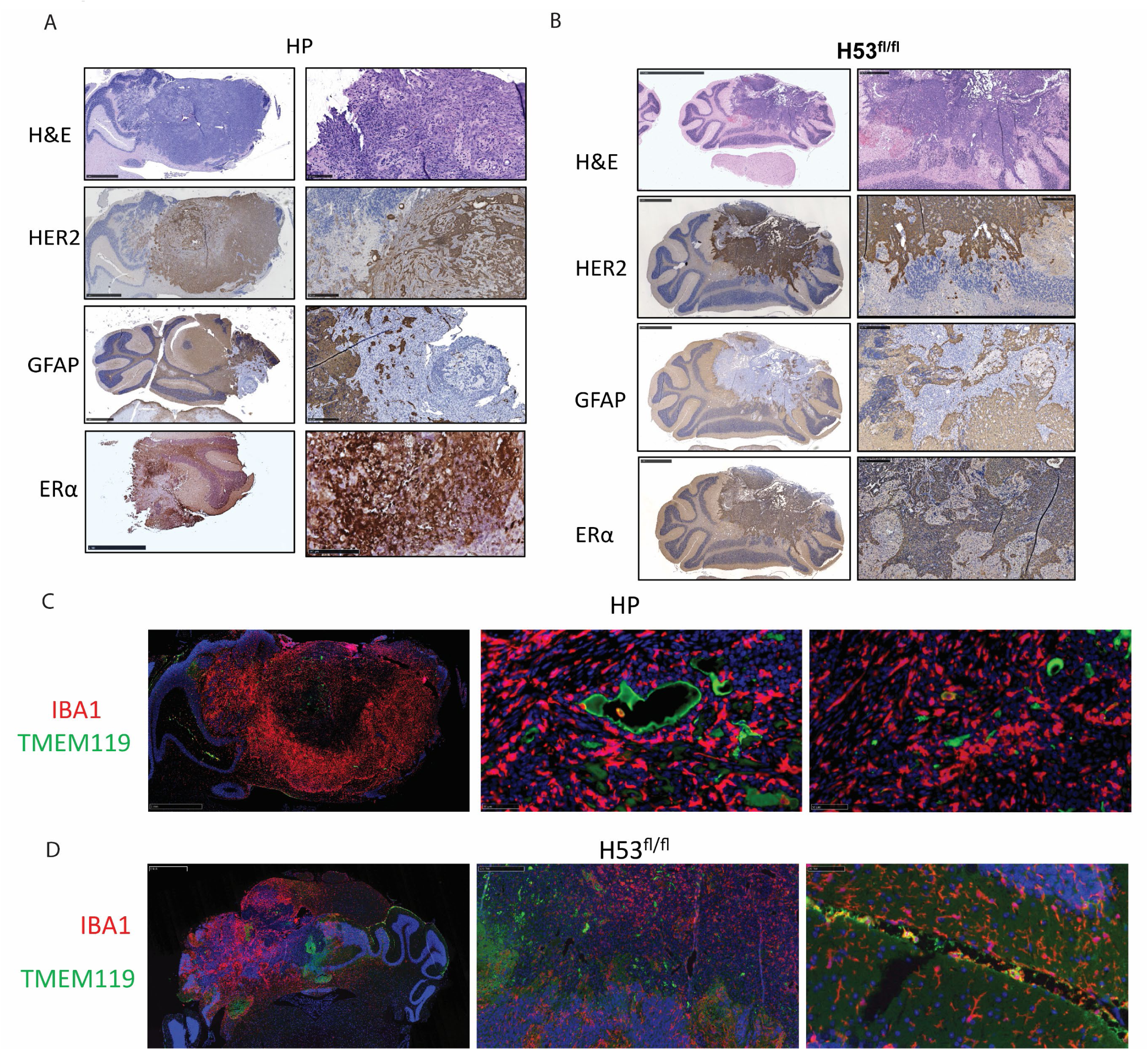
Characterization of cerebellum metastasis mouse model. (A) H&E and IHC stained with GFAP, HER2 and ER**α** antibodies in HP tumor organoid cells cerebellum metastasis model. Scale bars of low-power images are 5 mm. Scale bars of high-power images are 500 µm. (B) H&E and IHC stained with GFAP, ER**α**, and HER2 antibodies in the H53fl/fl tumor organoid cells cerebellum metastasis model. Scale bars of low-power images are 5 mm. Scale bars of high-power images are 500 µm. (C) IF stained with IBA1 and TMEM119 antibodies in HP organoid cells cerebellum metastasis model. Scale bars of low-power images are 5 mm. Scale bars of high-power images are 500 µm. (D) IF stained with IBA1 and TMEM119 antibodies in the H53fl/fl tumor organoid cells cerebellum metastasis model. Scale bars of low-power images are 5 mm. Scale bars of high-power images are 500 µm.

To further characterize the IBA1+ cells in these BCBM, we performed dual immunofluorescence to both IBA1 and the microglia marker, TMEM119 (Figure 2C-D). We observed very little co-localization of IBA1 and TMEM119, suggesting that the IBA1+ cells may be bone marrow-derived macrophages and not microglia. However, we acknowledge that a recent study reported that TMEM119 expression is downregulated in a subset of brain metastasis-associated microglia,^24^ therefore, further evidence, such as transcriptomics data, is needed to determine the identity of the IBA1+ cells. We checked whether the murine organoids that were injected contained any IBA1+ cells and we confirmed that the organoid cells injected to form these BCBMs did not contain IBA1+ cells (Figure S1). For the remainder of our experiments, we chose to focus on BCBM generated with the murine HP organoids.

### Near-infrared trastuzumab optical imaging of brain metastases

The BBB normally limits the penetration of many drugs and antibodies into the CNS. However, the BBB around brain metastases is abnormal and leaky, and has been termed the blood-tumor barrier (BTB).^5^ We had previously imaged lung metastases and mammary tumors in HP mice with optical imaging using a near-infrared fluorophore (NIR) conjugated to trastuzumab.^20^ To determine if NIR-trastuzumab imaging was effective for these cerebellar BCBM, we injected NIR-labeled trastuzumab into the tail vein of five cerebellar BCBM bearing mice. MRI images in Figure 3A confirm the presence of the cerebellar BCBM, and bioluminescence imaging (Figure 3B) also demonstrated the presence of cerebellar BCBM. 24 hours post NIR-trastuzumab injection, mice were shaved and underwent *in vivo* imaging (Figure 3C). NIR signal was readily detected through the intact skin and skull of all five mice. NIR-trastuzumab has an advantage over bioluminescence imaging as it can be performed *ex vivo* on dissected or even on formalin-fixed tissues (Figure 3D-E). This *ex vivo* signal is present because, unlike bioluminescence imaging, NIR imaging does not require ATP or living cells.^25^ Therefore, NIR imaging can be used to facilitate brain dissection. However, NIR signal is lost when the tissue is exposed to organic solvents used in paraffin embedding and routine histology (data not shown).

**Figure 3.**
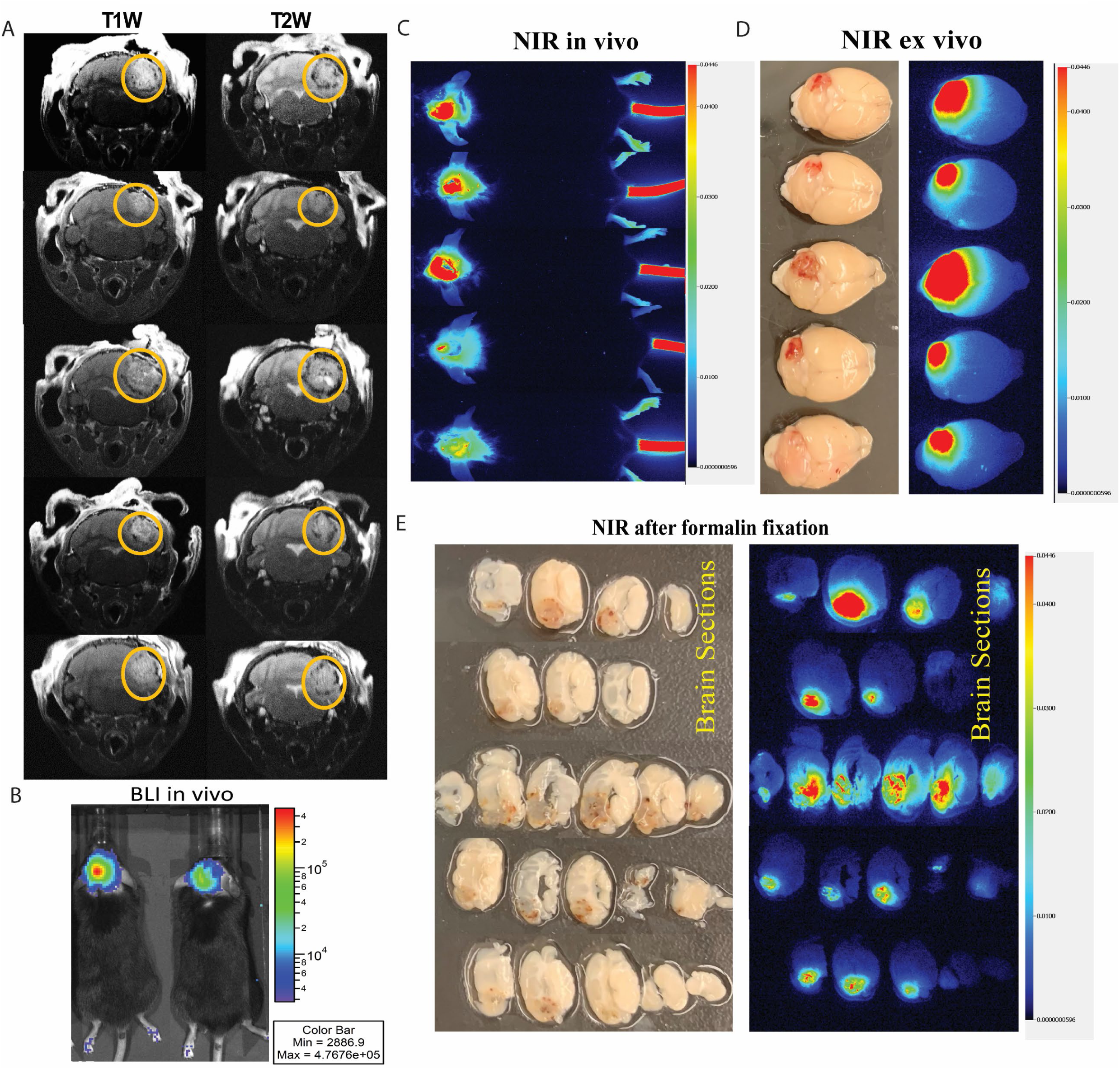
Optical imaging of brain metastases. (A) T1-weighted and T2-weighted MRI of stereotaxic injection at three weeks post-transplantation of HP organoids in C57BL/6 syngeneic mice. (B) Representative BLI images of the cerebellum tumor at 3 weeks post organoid cell transplantation of HER2+ tumor organoids. (C) Imaging of near-infrared fluorophore-labeled trastuzumab (NIR-trastuzumab) at 24 hours post-injection on the whole body of HP BCBM mice. (D) Left: Brightfield images of dissected and post-fixed cerebellum tumor at three weeks post-transplantation. Right: Imaging of near-infrared fluorophore-labeled trastuzumab (NIR-trastuzumab) at 24 hours post-injection on the whole brain of HP BCBM mice. (E) Left: Brightfield images of dissected and post-fixed cerebellum tumor slices at three weeks post-transplantation. Right: Imaging of near-infrared fluorophore-labeled trastuzumab (NIR-trastuzumab) at 24 hours post-injection on the brain section after formalin fixation derived from HP BCBM mice.

Overall, NIR-trastuzumab tumor-specific imaging is a potent tool for studying HER2+ BCBM. It demonstrates that monoclonal antibodies and conjugated antibodies can penetrate into these cerebellar BCBMs. Further, labeling trastuzumab with other imaging agents, such as PET isotopes like ^89^Zr or ^64^Cu, can provide a wide range of options for imaging HER2+ BCBMs.^26^ **Spatial transcriptomics reveals the role of Arg1 in the invasion clusters**

The tumor microenvironment (TME) plays a crucial role in the development, progression, and response to therapy in HER2+ breast cancer. The TME of HER2+ breast tumors is normally characterized by a complex interplay of various cell types, including cancer cells, stromal cells, immune cells, endothelial cells, and extracellular matrix components. HER2+ breast tumors often exhibit increased immune infiltration, with higher tumor-infiltrating lymphocytes (TILs) levels than HER2-negative tumors.^27^ Therefore, we investigated how the HER2+ BCBM tumor interacts with the unique TME in the cerebellum. We generated spatial transcriptomics data from two biological replicates of the cerebellar BCBM generated from HP organoids. After QC pre-filtration, qTune/qPlot, batch effect, and harmonizing the samples and clusters, we classified the samples into 9 clusters (Figure 4A, Table S1). The 9 clusters of the cells were further categorized as normal cerebellum, tumor cluster, and invasion cluster. In particular, the invasion cluster was differentiated into Cluster 1, which faced the normal cerebellum, and Cluster 6, which bordered the core of the tumor lesion, to help us better understand the progression of tumor invasion (Figure 4A). The UMI count distinguished between the HER2+ BCBM tumor and the normal area (Figure 4B), corresponding with the histological morphology (Figure 4C). The volcano plot highlighted the genes that were differentially expressed among tumor, invasion and normal cerebellum clusters (Figure 4D, Table S2). Specifically, we set a comparison group between Cluster 1, the invasive region adjacent to the normal cerebellum, and Cluster 6, the region including both the invasive perimeter and the tumor lesion core, to differentiate the gradual occurrence of metastasis (Figure 4D). Compared to the normal cerebellum and invasive clusters, many epithelial markers and inflammatory-associated genes were upregulated in the tumor clusters, including *Mmp3*, *Epcam*, *Krt7*, *Igha*, *Ighg2b*, *Igkc*, and *Anax1* (Figure 4D). In addition, some monocarboxylate transports like *Slc5a8* and cell-cell adhesion markers like *Col14a1*, *Fbln7*, and *Lum* were highly expressed in tumor clusters (Figure 4D). Interestingly, *Ltf*, which is a protein product found in the secondary granules of neutrophils, was upregulated in tumor populations. Some brain-dominant expressed genes like *Clip1* and *Clip2* were also found in the metastatic tumors, highlighting the unique molecular heterogeneity of the tumor cluster profile (Figure 4D). The invasion clusters exhibited enhanced gene signatures associated with cell-cell adhesion (*Col5a1*, *Col6a3*, *Col8a1*), angiogenesis (*Hmox1*), phagocytosis (*Arg1*), and metastasis (*Mmp12*, *Mmp13*) (Figure 4D). Importantly, *Arg1* gene expression were consistently upregulated in the invasion clusters across the replicate samples. As a marker of M2 macrophages, Arg1 had a role in immunosuppression and T cell activation^28^.

**Figure 4.**
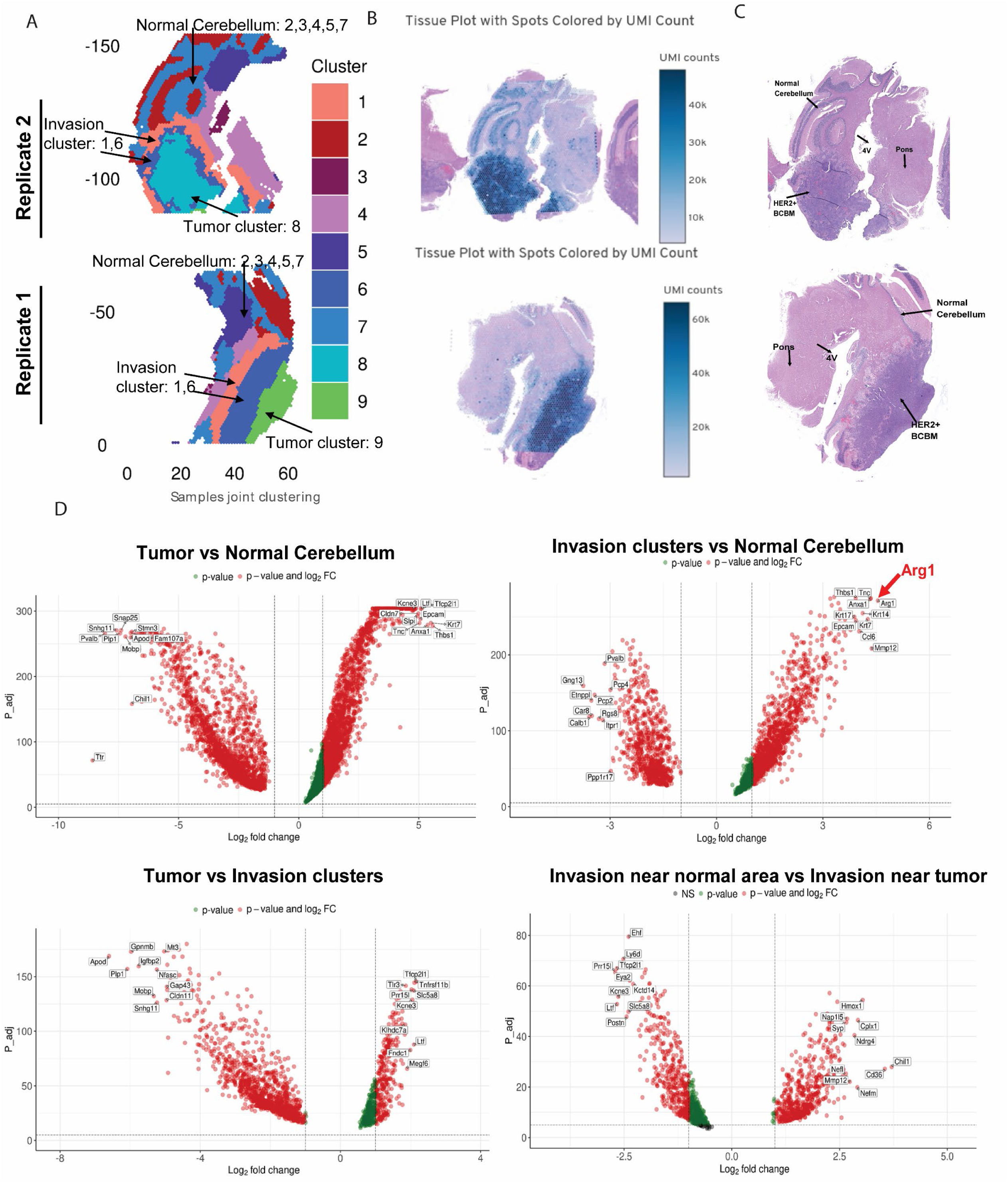
Spatial transcriptomics uncover a change in gene signature as HER2+ breast cancer invades the brain. (A) Sample joint clustering in the duplicate samples of spatial transcriptomics in the HER2+ breast cancer cerebellum metastasis model. The average expression of whole genome in each cluster is provided in Table S1. (B) Tissue plot with spots colored by UMI count in the duplicate samples. (C) H&E staining of the biological duplicate samples for spatial genome-wide sequencing. (D) Volcano plot illustrating genes meeting cutoffs for differential expression [log fold change (logFC2) >1, Padj. < 0.05] between Tumor versus normal cerebellum, Tumor versus invasion clusters, invasion clusters versus normal cerebellum, and invasion clusters near normal area versus invasion near the tumor. A list of the significantly altered genes in replicate two is provided in Table S2.

These results demonstrate that distinct gene expression panels exist in the tumor, normal cerebellum, and invasion clusters. *Arg1* in the invasion clusters suggests its role in pro-metastatic potency. However, it is unknown which cell types mediate the invasion by intersecting with normal cells in the cerebellum.

### Distinguished enrichment of pathways and cell types in the spatial landscape

HER2+ breast tumors can attract immune cells to the TME through various mechanisms, including the secretion of chemokines and cytokines. Tumor-infiltrating immune cells, such as T, B, natural killer (NK), and dendritic cells, interact with HER2+ cancer cells and influence tumor growth, invasion, and metastasis.^29^ We asked if HER2 activation in cancer cells can produce cytokines, chemokines, and growth factors that modulate the TME in the scenario of cerebellar metastasis. The GSEA analysis using the Hallmarks gene set indicated that tumor versus normal cerebellum differentially expressed genes (DEGs) were significantly enriched for genes related to inflammation, EMT, cell cycle, and metabolism (Figure 5A, 5B; Table S3). These data confirmed the distinct cluster properties in the tumor, normal cerebellum, and invasion edge.

**Figure 5.**
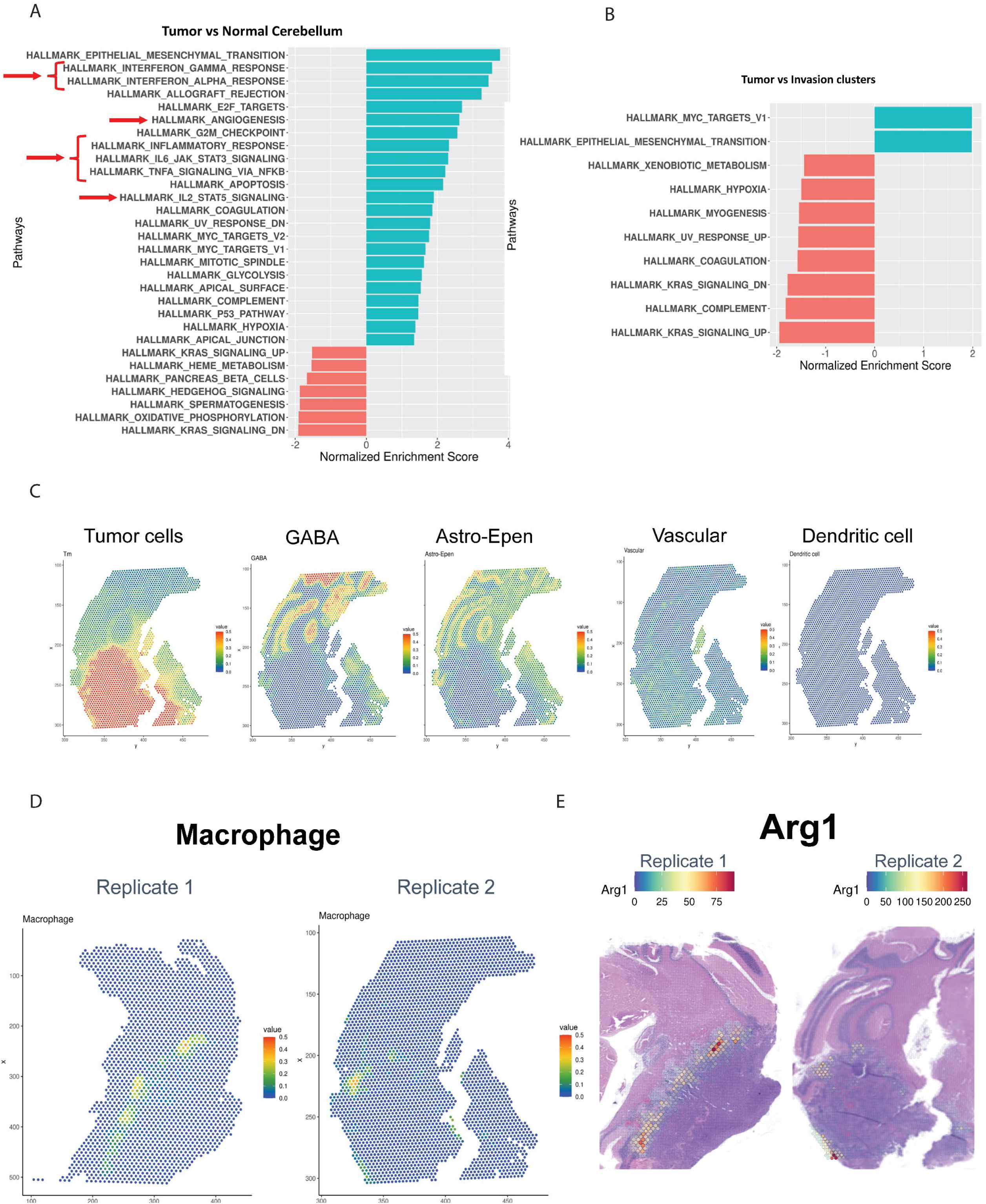
Cell type differentiation in tumor, invading, and normal cerebellum clusters. (A) Gene sets significantly enriched in tumors compared with normal cerebellum as identified by GSEA (p < 0.05)). ES (enrichment score) and-log10(p-values) of pathways are shown. GSEA was performed using the Hallmark gene sets in the Molecular Signatures Database (v7.5.1). A list of the hallmark genes in replicate one is provided in Table S3. (B) Gene sets significantly enriched in tumors compared with invasion clusters as identified by GSEA (p < 0.05). ES (enrichment score) and-log10(p-values) of pathways are shown from GSEA performed using Hallmark Gene sets in the Molecular Signatures Database (v7.5.1). A list of the hallmark genes in replicate one is provided in Table S3. (C) Integrated spatial distribution panel of cell types prediction in duplicate samples. Cell types are identified by a combination of snRNA and spatial transcriptomics technology. (D) Spatial distribution of macrophages as indicated in duplicate samples. D) Spatial distribution of Arg1 as indicated in duplicate samples.

The TME plays a critical role in shaping the behavior and response to therapy in HER2+ BCBMs. Elucidating the complex interactions between HER2 signaling and the TME components is essential for developing novel therapeutic approaches to overcome treatment resistance and improve outcomes for patients with HER2+ BCBMs. We next differentiated the cell type by integrating our spatial transcriptomics data with scRNA-seq data (Figure 5C). Tumor cell signature can be clearly seen by the red color in Figure 5C, most left panel. The GABA (neurons) and Astro-Epen-positive (glial) cell populations are mainly located in the normal cerebellum area. There are immune cells, myeloid leukocytes, and stroma cell types enriched in the tumor cohort; however, we did not see a significant difference in the infiltration of T cells, B cells, and dendritic cells. Interestingly, macrophage cells are significantly upregulated in the invasion clusters (Figure 5D), consistent with the *Arg1*spatial distribution (Figure 5E).

In summary, we identified macrophages at the edge of invasion, overlapping with increased Arg1 transcript expression seen in the spatial transcriptomics data. Understanding the interactions between HER2+ BCBM and the TME in the cerebellum is essential for developing effective therapeutic strategies for brain metastasis.

### Spatial transcriptomics panels reveal cluster-type “gradient” as HER2+ tumor invades the cerebellum

Previous studies have shown that ErbB2 and ErbB4 are constitutively expressed in granule cells during the first 2-3 weeks postnatally,^30^ suggesting that dimerization such as ErbB2/ErbB4 or ErbB4/ErbB4 may be important for processes such as granule cell migration, neuronal connection, and synaptic formation. We analyzed the gene expression in specific cell subtypes. *HER2*, *EGFR*, and *HER3* were upregulated in the tumor clusters along with increased downstream signaling genes, *Akt* and *Mapk1*, suggesting that a combination of HER2 with EGFR or HER3 was required to form functional receptors in cerebellar metastatic tumors (Figure 6A). In contrast, *HER4* is mainly expressed in the normal cerebellum clusters, revealing its role in the normal cerebellum (Figure 6A).

**Figure 6.**
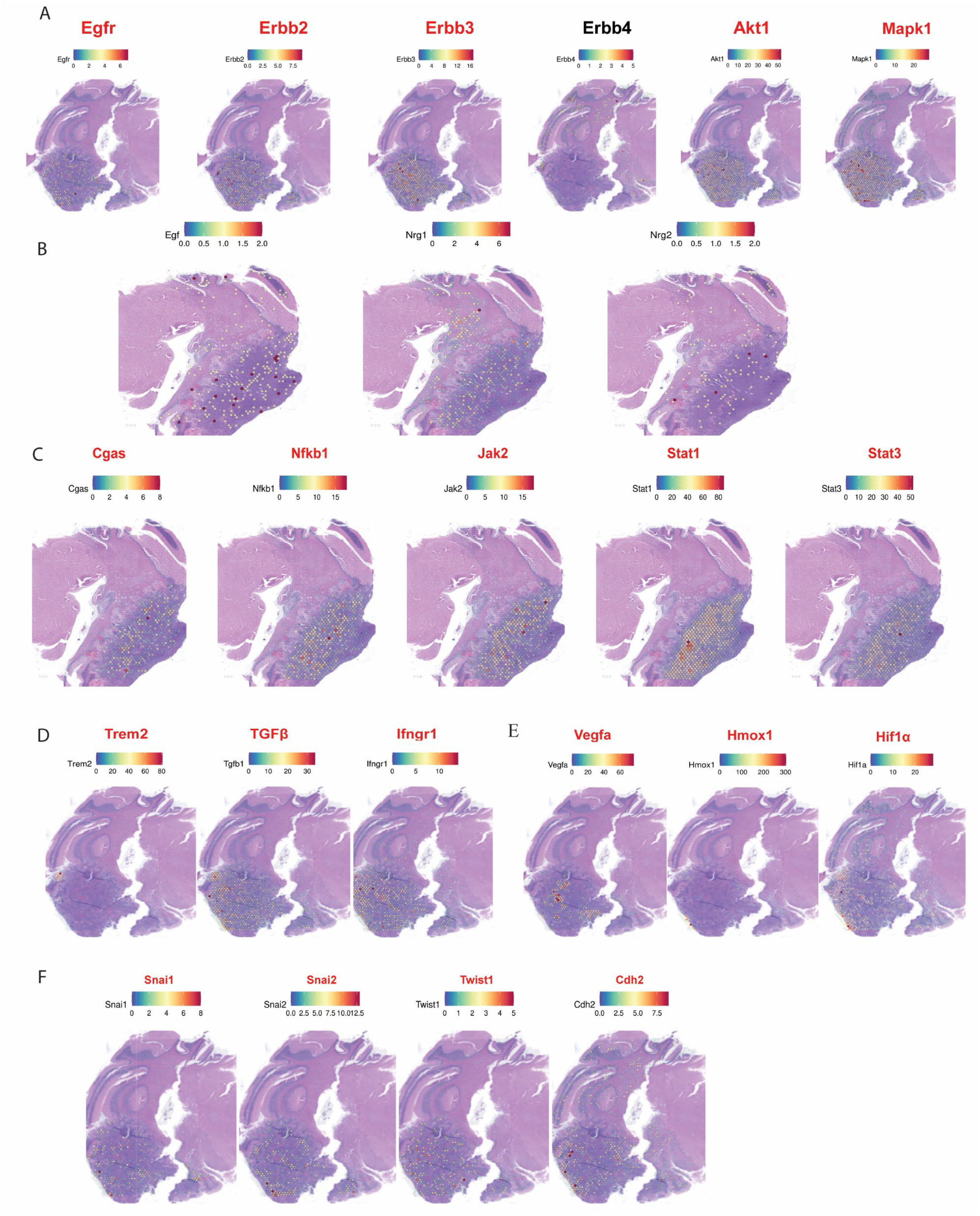
Inflammation and macrophage cells are drivers in the invading edge. (A) Spatial feature plot of key genes and factors related to HER2 signaling in the representative tissue sample. (B) Spatial feature plot of HER2 ligands in the representative tissue sample. (C) Spatial feature plot of key genes or transcript factors regulating inflammation signaling in the representative tissue sample. (D) Spatial feature plot of macrophage activators in the representative tissue sample. (E) Spatial feature plot of key genes or transcript factors regulating angiogenesis in the representative tissue sample. (F) Spatial feature plot of key genes or transcript factors regulating EMT (epithelial-mesenchymal transition) signaling in the representative tissue sample.

The EGF family ligands, *Egf*, *Nrg1*, and *Nrg2* are expressed in both the normal cerebellum and the BCBM tumor (Figure 6B). Consistent with the GSEA results (Figure 5A), a number of inflammatory genes are increased in the BCBM tumors, including *Cgas*, *Jak2*, *Stat1*, *Stat3* and *Nfkb1* (Figure 6C). High expression of *Trem2*, *TGFβ*, and *Ifngr1* in Figure 6D suggests that HER2+ BCBM attract immunosuppressive cells like regulatory T cells (Tregs), myeloid-derived suppressor cells (MDSCs), and tumor-associated macrophages (TAMs), thereby creating an immunosuppressive TME. HER2 signaling can upregulate cytokines such as IL-6 and chemokines like CCL2, promoting a pro-inflammatory and immunosuppressive environment conducive to tumor growth and metastasis. In fact, inflammation is the key step that will result in neurodegeneration disease^31^ and primary brain tumors.^32^ Cerebellar granule cells undergo major processes of neural development and synaptic connections, which are sensitive to the chemokines and cytokines secreted by immunogenic or tumor cells. *Vegfa*, *Hif1a*, and *Hmox1* are highly expressed in the invasion clusters correlating with angiogenesis’s maturation (Figure 6E).

Particularly, Tgfb1 expression levels are much higher than other cytokines (Figure 6D). In addition, Ifngr, a receptor of type II class of interferons, which is Jak-Stat-Gas pathway-specific cytokines, is mainly localized in the tumor area (Figure 6D). Both cytokines and growth factors can trigger Jak-Stat-Gas signaling. Sting pathway detects extracellular DNA, which can indicate either a foreign invader, such as a virus, or damage to host tissue or cells. HER2 recruits AKT1 to disrupt STING signaling and suppress antiviral defense and antitumor immunity.^33^ Consistent with this, *Sting1*, *Ifna*, and *Ifnb1* are not expressed in both normal cerebellum and BCBM cohort (data not shown), suggesting Sting may be a runaway gap for inflammation mediated by Ifngr-induced Jak-Stat-Gas pathway in HER2 positive breast cancer cerebellum metastasis.

Epithelial mesenchymal transition (EMT) is the mechanism for invasion in many cancer types. *Snail*, *Snail2*, *Twist1* and *Cdh2* are increased in both tumor and invasion clusters (Figure 6F), indicating the EMT transition is a necessary step for the cerebellum metastasis. HER2 is reported to induce TGFβ/SMAD signaling to further activate the SNAIL and SNAIL2,^34^ this mechanism matches with the Erbb2-Tgfb1-Snail/Snail2/Twist1 cascade we observed.

In summary, inflammation in the tumor microenvironment of the HER2+ BCBM has a potential role in tumor development. Understanding the complex interactions between HER2 signaling and the immune system is crucial for developing effective immunotherapeutic strategies and improving outcomes for patients with HER2+ BCBMs.

### IL34 secretion by breast cancer cells induces ARG1+ macrophages, and a blocking antibody reduces BCBM growth

We hypothesized that cytokines or chemokines specifically secreted by the tumor organoid cells could induce macrophage polarization at the edge of invasion in the BCBM models. The spatial transcriptomics data showed that Il34 was highly expressed in the tumor area (Figure 7A).

**Figure 7.**
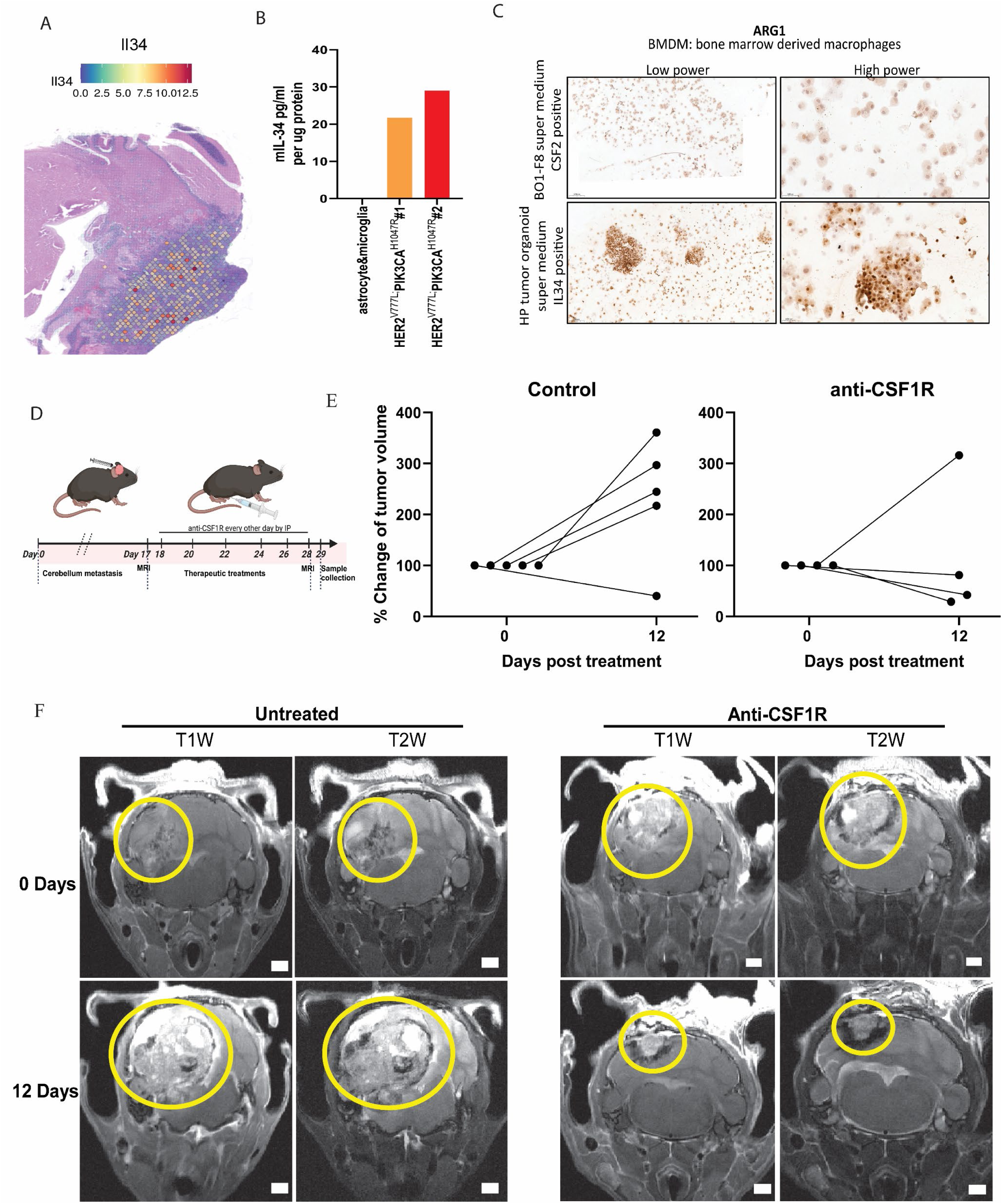
IL34-induced Arg1 expression at the edge of invasion in cerebellum metastasis of HER2+ breast cancer. (A) Spatial feature plot of Il34 in the representative tissue sample. (B) ELISA test result of IL34 protein level from astrocytes and microglial cells, HER2*^V777L^*; PIK3CA*^H1047R^* tumor organoid cells. (C) The mouse BMDM was cultured with a conditioned media supernatant from BO1^39^ cells or HP tumor organoid cells for 24 hours, and the IHC staining of ARG1 was performed on cytospin slides of the co-cultured cells. (D) Schematic representation of anti-CSF1R mAb treatment in cerebellar tumors of BCBM mouse model. (E) percent change of tumor volume in the control and anti-CSF1R mAb groups. (F) The representative T1W and T2W images of MRI showed the cerebellar tumor volume change at 0 days and 12 days post-treatment in control and anti-CSF1R mAb groups. 5000 cells of HER2*^V777L^*; PIK3CA*^H1047R^* tumor organoid were injected into the mouse cerebellum for modeling as described in the method.

Cerebellar neurons do not express IL-34, and IL-34 mutant animals do not change the microglia cell number in the cerebellum and brainstem among the whole brain regions.^35^ Therefore, we tested whether the breast cancer cells expressed IL34 protein using an ELISA assay. IL34 was detected in protein lysates from HP breast cancer cells but not from C57BL/6 mouse brain astrocytes or microglia (Figure 7B). Bone marrow-derived macrophages from C57BL/6 mice were incubated with conditioned media from the HP breast cancer cells, and Arg1 protein expression was induced (Figure 7C).

Cerebellar microglia homeostasis requires CSF1R, but they are not Il34 dependant.^36^ Microglial cells have shown distinct transcriptional profiling compared with other cell types in the mouse cerebellum.^37^ We wanted to determine how macrophages and microglia are differentially localized and function in the tumor and invasion clusters of our BCBMs model (Figure S2A; Figure 2C; Figure 5E). We first checked for macrophage markers in the tumor area (Figure S2A). The differential expression of ARG1 was found at the edge of the invasion area near the normal cerebellum clusters (Figure 4D). Macrophage enrichment is a hallmark of pancreatic cancer, and Arg1-expressing macrophages are a key driver of immune suppression.^38^ Consistent with results in Figures 4D and 5E, Arg1 is specifically expressed in the invasion cell populations which is distinct from the spatial expression of Itgam (Cd11b), Iba1, Tmem119, Creb1, Cd68, Ptprc(Cd45) seen mainly in the tumor clusters (Figure S2A). Intriguingly, Camk2a, which is a functional marker, is predominantly expressed in the normal brain microglia cells (Figure S2A). Chemokines and cytokines that regulate macrophages like Il4, Ccr2, and Ccl2 are upregulated in the tumor and invading cohort of brain metastasis (Figure S2B). These results suggest that Il34 can induce Arg1-positive macrophage cells in the invasion border of the HER2+ BCBM model.

We confirmed ARG1 and IL34 expression at the edge of invasion in the BCBM models (Figure S3A, S3B). Additionally, there are some ARG1 and IL34 positive cells in the HP primary breast tumor and HP breast cancer metastatic lung tumor but not at the border of the tumor invasion area. Mouse bone marrow-derived macrophages (BMDM) were cultured with conditioned media supernatant harvested from BO1^39^ cells or HP tumor organoid cells after 24 hours in culture. We found that the conditioned media supernatant from the HP breast tumor organoids can polarize the macrophage cell morphology (Figure S3C). BMDM and mouse BV2 cells were cultured with conditioned media supernatant from BO1^39^ cells or HP tumor organoid cells for 24 hours, with slides for IHC produced by cytospin, IHC staining for ARG1 (Figure 7C, S3D) and IL34 (Figure S3E) was performed on the cytospin slides. ARG1+ cells were observed in both BMDM and BV2^40^ cells after coculture with HP breast tumor organoid-conditioned media supernatant for 24 hours (Figure 7C, S3D). We also detected the expression of IL34 in the BV2 cells co-cultured with HP breast tumor organoid conditioned media supernatant (Figure S3E).

CSF1R is the receptor for IL34, and a blocking antibody to mouse CSF1R is available.^41, 42^ We already demonstrated that monoclonal antibodies can penetrate *in vivo* into these cerebellar BCBM (Figure 3). To model the treatment of brain metastases, we treated mice with established cerebellar BCBM with this CSF1R blocking antibody, and we observed BCBM shrinkage in 3 out of 4 (75%) mice (Figure 7D-F, S4A, S4B). This was a short-term experiment with a 12-day duration of antibody treatment (timeline shown in Fig. 7D). In contrast, control mice showed dramatic tumor growth over the same time period (Figure 7E, S4B). MRI images of representative mice show BCBM shrinkage with CSF1R antibody treatment (Figure 7F) and BCBM growth in the control group (Figure 7F).

In conclusion, Il34 secretion from HER2+ BCBM can induce border-associated Arg1-positive macrophages. Blocking the Il34-CSF1R by anti-CSF1R mAb can be a potentially effective immunotherapy in treating brain metastases tumors and could be combined with existing treatments for HER2+ brain metastases.

## DISCUSSION

In this paper, we generated a novel, immunocompetent animal model for cerebellar BCBM using transplantation of breast cancer organoids from one B6 mouse to another B6 mouse. We identified an important role for Arg1+ macrophages at the tumor periphery, and we demonstrated that inhibiting these macrophages with a CSF1R-blocking antibody produced tumor shrinkage. While there have been considerable advances in the treatment of HER2+ BCBM, including the antibody-drug conjugate (ADC) trastuzumab deruxtecan,^14, 43^ the tyrosine kinase inhibitor tucatinib,^44^ and focused radiation therapy techniques,^45^ most patients with HER2+ BCBM will die a neurological death from progressive brain metastasis.^43^ Therefore, new strategies and additional approaches to treat brain metastases are needed. Our results with the CSF1R blocking antibody potentially provide such a new approach.

Inflammation is one of the hallmarks of cancer,^46^ and numerous studies have examined the role of tumor-associated macrophages.^47^ Attempts to inhibit or deplete tumor-associated macrophages with CSF1R tyrosine kinase inhibitors such as BLZ945 or JNJ-40346527 have unfortunately been unsuccessful,^48, 49^ possibly because of activation of parallel or redundant pathways or off-target effects on closely related tyrosine kinases.^50^ Targeting CSF1R with a monoclonal antibody has a number of potential advantages. First, antibody drugs often have greater affinity and specificity for the target protein.^51^ Second, monoclonal antibodies typically have a much longer half-life than small molecule kinase inhibitors, thereby providing sustained target inhibition.^52^ Third, immunological effects of antibody binding such as antibody-dependent cellular cytotoxicity (ADCC) and antibody-dependent cellular phagocytosis (ADCP) provide additional mechanisms for drug action beyond the kinase activity inhibition produced by small-molecule kinase inhibitors.^51^ The recent FDA approval of axatilimab for the treatment of another cancer-related condition, chronic graft versus host disease,^13^ offers a new class of targeted therapies for CSF1R and tumor-associated macrophages.

### Limitations of this study

We acknowledge that direct injection of breast cancer cells into the cerebellum bypasses the intravasation and extravasation steps in the metastatic cascade. However, direct injection is the most experimentally tractable method to study cerebellum metastasis, and it provided highly useful information about the changing microenvironment around a growing metastasis. We also acknowledge that this study does not compare cerebellum metastasis to other brain regions. We plan to perform this comparison in our next study. Further, no human samples were analyzed in this study, and we also plan to include human BCBM neurosurgical samples in future studies. Finally, spatial transcriptomics platforms and methods are rapidly evolving. In late 2024, 10x Genomics released the Visium HD platform, which provides up to 2 μm resolution.^53^ The availability of single-cell or near-single-cell spatial techniques will provide further understanding of brain metastases. Nevertheless, despite these limitations, this study provides valuable information about how inflammation affects cancer invasion in the cerebellar metastasis of HER2+ breast cancer and identified the role of Arg1+ macrophages at the invading edge of the HER2+ BCBM.

## Conclusions

We developed a novel, immunocompetent animal model of cerebellar brain metastases from HER2+ breast cancer and used it to identify a key role for IL34-induced Arg1+ macrophages.

Inhibiting IL34-CSF1R signaling with a CSF1R-blocking antibody produced tumor shrinkage. Future studies will examine if IL34-induced Arg1+ macrophages play a similar role in HER2+ BCBM in other brain regions and if this mechanism also contributes to the growth of brain metastases from other types of cancer.

## Graphical summary

Cheng et al. demonstrate that there is a vicious cycle between the breast cancer cell and Arg1+ macrophages in cerebellar brain metastasis. Breast cancer cells secrete IL34, which binds CSF1R on the macrophages, stimulating inflammation and tissue damage, which promotes further cancer invasion and growth. CSF1R blocking antibody interrupts this cycle and produces metastasis shrinkage.

## Resource availability

Spatial transcriptomics data generated in this study will deposited in the Gene-Expression Omnibus with an accession code once published. All other data supporting the findings of this study are available from the corresponding author upon reasonable request. Source data are provided in this paper. All the analyses were based on standard algorithms described in the Methods and referenced accordingly. Custom algorithms are to be made available upon request. All other raw data are available upon request from the corresponding author.

## Acknowledgments

We thank Dr. Katherine Weilbaecher’s lab for the help with BMDM isolation and BO1 condition medium. We also thank Dr. Gilbert Gallardo’s lab for the help with the isolation of mouse primary astrocyte cells. We thank Drs. Max Wattenberg and Sonika Dahiya for helpful discussions. This work was supported by the Department of Defense Breast Cancer Research Program grant number BC170330 (to R.B.); the Ohana Breast Cancer Research Fund and the Foundation for Barnes-Jewish Hospital (to R.B.); and NIH grants F30 CA268778 (to M.S.M.), S10 OD027042 and S10 OD025264 (to the Mallinckrodt Institute of Radiology, Molecular Imaging Center) and P30 CA091842 (to the Siteman Cancer Center Small Animal Cancer Imaging shared resource).

## METHODS AND RESOURCES

### Mice

All animal studies were approved by the Institutional Animal Care and Use Committee (IACUC) at Washington University in St. Louis. HER2^V777L^ transgenic mice were generated using TALEN-based genome editing as previously described.^19, 20^ *PIK3CA*^H1047R^ mice were purchased from The Jackson Laboratory, strain #:016977. *Trp53*^fl/fl^ mice were purchased from The Jackson Laboratory, strain #: 008462. All animal experiments were performed under the Institutional Animal Care & Use Committee (IACUC) approved protocols.

### Establishment of mammary gland organoids from murine breast cancers

Mammary gland tumors were resected from tumor-burdened HP (HER2V777L; PIK3CAH1047R) or H53(HER2V777L; Trp53fl/fl) mice. The tumors were minced and tumor clusters were isolated by a modification of Ewald’s protocol.^54^ In brief, the minced breast tumors were enzymatically dissociated at 37 degrees C in collagenase solution for organoids^54^ with agitation using a Gentle MACS Octo Dissociator with heaters and the program, 37C_m_LDK_1, run twice. Dissociated tissues were centrifuged at 520g for 10 minutes at (22 degrees C). The supernatant was discarded and the pellet was washed with Ad DME/F12, supplemented with 1X Glutamax, 1X HEPES and 1X penicillin/streptomycin (+++ media) and centrifuged at 520g for 10 minutes at 22 degrees C. The cell pellet was treated with 1 ml of 1X RBC lysis buffer (Invitrogen-eBioscience) for 5 minutes followed by neutralization with 40ml PBS. After centrifugation, the cell pellet was treated with 4ml of 4units/ml of DNAse-I in +++ media, with shaking for 5 minutes. After DNAseI treatment, 6ml of +++ media was added, and the tube was centrifuged at 520g for 5 minutes. The cell pellet was washed in 10ml +++ media, and after centrifugation, the pellet was resuspended in 90% Matrigel GFR and seeded in a 6-well ultra-low adhesion plate (Corning 3471). The embedded cells were overlaid with 3 mL/well mouse breast tumor medium^55^ (advanced DMEM/F12 supplemented with penicillin/streptomycin, 10 mmol/L HEPES, Glutamax, 1 × B27 [all from Thermo Fisher Scientific], 0.5X primocin (Invivogen), 125 μmol/L N-acetyl-cysteine (Sigma), 50 ng/mL human epidermal growth factor (rhEGF, Peprotech)), and 10% Rspo1-Noggin–conditioned medium. (The Noggin and R-spondin combined expression plasmid to generate conditioned media in the 293T cell line was a gift from Blair Madison, PhD and Anil Rustgi, MD). The mouse breast tumor medium was supplemented with the Rho Kinase inhibitor, Y-27632, at 10uM for the first feeding after splitting. The growth medium was replaced every 3-4 days, and organoids were passaged every 7-10 days by treatment with Cell Recovery Solution (Corning), followed by digestion with TrypLE Express and mechanical dissociation and re-embedding in fresh Matrigel and ultra-low adhesion plates.

### Luciferase labeling

Organoid cells were transduced with a lentivirus-LUC luciferase reporter. Cloning selection was performed to isolate stable clones with luciferase expression. Luciferase labeling was confirmed using quantification of *in vitro* bioluminescence signal.

### Stereotactic intracranial injections

Mice were anesthetized using 2% isoflurane with 100% oxygen for the duration of the operation. Prior to an incision above the base of the skull, the skin was sterilized with three alternating wipes of ethanol and betadine. Injections were done with a 28G needle with a standard 12° beveled needle point mounted on mouse stereotactic frame courtesy of the Albert Kim Lab. After sterilization, the skull was exposed, and a small burr hole was drilled using a dental drill at the coordinates of the desired injection site according to the Paxinos and Franklin’s mouse brain atlas, 5^th^ edition. A microsyringe pump was used to deliver 3uL of organoids, which had been dissociated into small clusters, into the cerebellum at a fixed rate of 0.4 μL/min. Following injection, the needle was left in place for an additional 5 minutes to allow the injection solution to diffuse before slowly retracting. The injection site was sterilized with ethanol before the drilled hole was sealed with Gorilla Glue. The incision site was sutured, and the animals were allowed to recover on a heating pad before being returned to the animal cage for post-operative monitoring.

Organoids were removed from Matrigel using Cell Recovery Solution (Corning), washed, digested in TrypLE Express, and resuspended in Dulbecco’s Modified Eagle Medium (DMEM). During the optimization process, different organoid cell numbers were injected into the cerebellum, ranging from 1000 cells to 5000 cells. The dissociated organoids for injection were prepared on the day of transplantation and kept on ice until the surgery was completed.

### Small animal MRI

Mice underwent MRI starting 3 weeks after stereotactic injection. Gadolinium-based contrast agent (GBCA) was injected intraperitoneally before imaging. While in the MRI scanner, the mice were anesthetized using isoflurane, and their heads were fixed in place using ear pins and bite bars. *In vivo* MRI studies were performed on a 9.4T MR scanner (Bruker Biospec, Ettlingen, Germany) using a 86mm diameter transmit coil and a 4-channel mouse brain cryo-probe surface coil. The animals were anesthetized with a mixture of isoflurane in oxygen (4% for 90s induction, 1% for maintenance), and the animal temperature was maintained at 37 ± 1 °C, using a circulating water bath and heating pad placed over the animal. Respiration and temperature was monitored during MRI acquisition with SA Instruments (SA Instruments, Inc. Stony Brook, NY, USA) sensors. Immediately prior to imaging, each mouse was given a 300mL i.p. injection of 2:10 Dotarem® (Guerbet) diluted in saline. The MRI protocol consisted of a T2 weighted RARE image (TE = 35 ms, TR = 2750 ms), and a T1 weighted RARE image (TE = 7.4 ms, TR = 800 ms). For each scan, 32 slices were acquired using a field of view of 16 x 16 mm2, an in-plane spatial resolution of 62.5 x 62.5 mm2 and a slice thickness of 500 mm. The total scanning time per animal was 20 minutes. Tumors were manually segmented using ITK Snap (http://www.itksnap.org/).

### H&E, IHC, and immunofluorescence staining

Tumor tissues were fixed with 10% formalin, embedded in paraffin, and cut into 5-mm sections (Digestive Diseases Research Core, Washington University School of Medicine). For H&E, slides were stained with hematoxylin, rinsed, and then counterstained with eosin, before being mounted and imaged. For IHC and IF, sections were deparaffinized, hydrated, and treated with heat-activated antigen unmasking solution (Vector Laboratories) before incubation overnight with primary antibody at 4 degrees C. For immunofluorescence staining, antibody binding was visualized with Alexa Fluor 488 or 555 or 647 fluorochromes and then counterstained with DAPI-containing mounting media (Sigma). For IHC staining, slides were treated with HRP conjugated anti-species secondary antibodies, followed by visualization with DAB substrate (Cell Signaling Technology) and then counterstained with hematoxylin (Thermo Fisher Scientific). Immunostaining was performed using the following primary and secondary antibodies. Primary antibodies include rabbit ERBB2 (CST, 2165S), mouse ERBB2 (CST, AF2967-SP), ERa Antibody (D-12) (Santa Cruz Biotechnology, SC-8005), GATA-3 Antibody (Santa Cruz Biotechnology, SC-268), Iba1/AIF4T (E404W) antibody (CST 17198S), TMEM119 antibody (CST 41134S), Arginase-1 antibody (CST 93668S), mIL-34 antibody (R&D systems AF5195). Secondary antibodies for IF include anti-rabbit IgG (H+L) Alexa Fluor 555 (Invitrogen, A-31572), anti-goat IgG (H+L) Alexa Fluor Plus 488 (Invitrogen, A32814), Goat anti guinea pig (H + L) FITC (Fitzgerald Industry International, 43R-1095). Secondary antibodies for IHC include SignalStain Boost IHC Detection Reagent (HRP, Rabbit) (Cell signaling technology, 8114S), SignalStain Boost IHC Detection Reagent (HRP, Mouse) (Cell signaling technology, 8125S) and anti-sheep IgG (HRP conjugated) (R&D Systems HAF016).

### *In vivo* CSF1R blocking antibody treatment

CSF1R blocking antibody was purchased from BioXCell (Clone AFS98, Catalog number BE0213). For this experiment, mice were stereotactically injected with 5000 tumor organoid cells. After 17 days, treatement was started with CSF1R blocking antibody injected intraperitoneally into the mouse every two days.^56^ The dosage on days 18,20,24,26 was 400**μ**g/mouse, while on days 22, 28 it was 100**μ**g/mouse.

### In vivo BLI

For BLI of live animals, mice were injected intraperitoneally with 150 μg /g D-luciferin (Gold Biotechnology, St. Louis, MO) in PBS, anesthetized with 2.5% isoflurane, and imaged with a charge-coupled device (CCD) camera-based system (IVIS 50, PerkinElmer, Waltham, MA; Living Image 4.3.1, exposure time 10-60 seconds, binning 8, field of view 12cm, f/stop 1, open filter, ventral view). Luminescence was displayed as photons/sed/cm2/sr. Regions of interest (ROI) were defined manually over the lower abdomen using Living Image 2.6 with measurements reported as photons/sec.

### Near-Infrared Optical Image

NIR-trastuzumab was injected into the tail vein of HP or H53^fl/fl^ organoid transplanted mice bearing breast tumor one day prior to imaging. The near-infrared optical imaging was taken using the Pearl Trilogy (LI-COR, Lincoln, NE) in the Washington University Molecular Imaging Center (MIC). The PearlCam and ImageStation software was used for the measurement and analysis.

### Visium spatial transcriptomics on mouse cerebellum brain metastasis

Tissue prepared for sectioning and staining start with RNA quality assessment. RNA quality assessment of FFPE tissue blocks (RNA extraction and QC) needs to meet the DV200 Quality Score = >30% to proceed with assay using Agilent BioAnalyzer or TapeStation Instruments. Approved slides thickness at 5 μm sections compatible with the Visium CytAssist. 20x resolution imaging on Zeiss Axio Scan 7 Brightfield / Fluorescence Slide Scanner was used for downstream Space Ranger Analysis. Sample/Slide was prepared with decrosslinking and library Construction on 6.5mmx 6.5mm capture area. Visium CytAssist Spatial Gene Expression kit was used for FFPE (Mouse Transcriptome, 6.5 mm, 4 rxns, PN-1000521,10x Genomics). The probe processes include probe hybridization/Wash, probe ligation/Wash, RNA digestion and tissue removal, probe extension and elution, preamplification and cleanup and Index PCR. CytAssist Brightfield images generated for downstream Space Ranger Analysis. For sequencing information, 100M total read pairs was targeted for each sample on the Illumina NovaSeqX Plus instrument (NovaSeq X Plus 10B at 300 cycle, 10M targeted 2×150 Clusters).

### Visium spatial data analysis

Initial visium spatial data analysis was performed using 10X Genomics Space Ranger software(v 2.1.1). Space Ranger includes two pipelines relevant to spatial gene expression experiments which are mkfastq and count. Spaceranger mkfastq wraps Illumina’s bcl2fastq to demultiplex the sequencing runs and to convert barcode and read data to FASTQ files. Space Ranger count then takes a bright field slide image and FASTQ files from mkfastq for alignment, tissue detection, fiducial detection, and barcode/UMI counting. After automated alignment and tissue identification, the spaceranger Count pipeline utilized the full-resolution slide image to prepare data for visualization within Loupe Browser (10xGenomics:https://support.10xgenomics.com/single-cell-gene-expression/software/visualization/latest/what-is-loupe-cell-browser). The region of interest of the tissue was identified and marked using Loupe Browser and subsequently selected for downstream analysis.

Output of the count pipeline contains expression matrix which was used for further analysis on the designated ROIs. For sample 1, the ROI consisted of 2,184 spots with a mean read depth of 30,825 reads per spot, while for sample 2, the ROI included 2,976 spots with a mean read depth of 28,271 reads per spot. Analysis was conducted using the Seurat(version 4.9.9.9086:Source: <u>vignettes/spatial_vignette.Rmd</u>) package in R. The quality control and subsequent analysis were performed on the spot-level expression data along with the associated tissue slice image produced by Space Ranger. During quality control step, the spots with number of unique molecules less than 500 and number of unique genes less than 10 were filtered out. After initial filtering, the final dataset retained 2,150 spots for sample1, while the spot count for sample2 remained unchanged. In Seurat, the filtered data is first normalized and scaled to account for variance in sequencing depth across data points. On normalized, scaled data, linear dimensionality reduction was performed by calculating 50 principal components using the most variably expressed genes in the spatial data. Spots were grouped into 9 clusters for de novo cell type discovery using BayesSpace’s qTune() and spatialCluster() functions and graph-based clustering approaches with visualization of spots was achieved through the use of clusterPlot().^57^

The Combined use of scRNA-seq and Spatial transcriptomics technologies on mouse brain specimen is expected to provide novel insights into the molecular, cellular and spatial organization of distinct breast cancer metastasis with that of the surrounding tumor microenvironment. Publicly available single cell datasets were used to identify the underlying composition of cell type (Allen Institute for Brain Science (2004). Allen Mouse Brain Atlas. Available from mouse.brain-map.org. Allen Institute for Brain Science). ^58, 59^ Reference based deconvolution workflow – Spacex-R ^60^ was applied to uncover cellular heterogeneity and spatial arrangement of cell types of the samples, thus providing for each spot a probabilistic classification for each of the scRNA seq derived classes. Post spot deconvolution, differentially expressed genes were identified using FindMarkers() from the Seurat package (Satija, R. Seurat-Analysis, visualization, and integration of spatial datasets.) for regions such as invasive region (near the tumor and near the normal area), Tumor and normal cerebellum. The top 10 upregulated and downregulated DEGs were visualized using a volcano plot (https://github.com/kevinblighe/EnhancedVolcano.). Then, gene set enrichment analysis was performed with above found DEGs using fgsea R package^61^ by utilizing the Molecular Signatures Database (MsigDB: Dolgalev I (2025). msigdbr: MSigDB Gene Sets for Multiple Organisms in a Tidy Data Format. R package version 2023.1.1, https://igordot.github.io/msigdbr/.). MSigDb has 50 predefined hallmark gene signatures which were leveraged to obtain the biological states or processes.

## Abbreviations

ADC: Antibody-drug conjugates.
ADCC: antibody-dependent cellular cytotoxicity
ADCP: antibody-dependent cellular phagocytosis
BBB: Brain blood barrier
BCBM: Breast cancer brain metastasis
BMDM: Bone marrow-derived macrophages
BTB: Brain tumor barrier
HER2: Human Epidermal Growth Factor Receptor 2
NIR: near-infrared
PDX: Patient-Derived Xenografts
UMI: unique molecular identifier
T-DXd: Trastuzumab deruxtecan
TME: Tumor Microenvironment.

---

**Figure S1.**
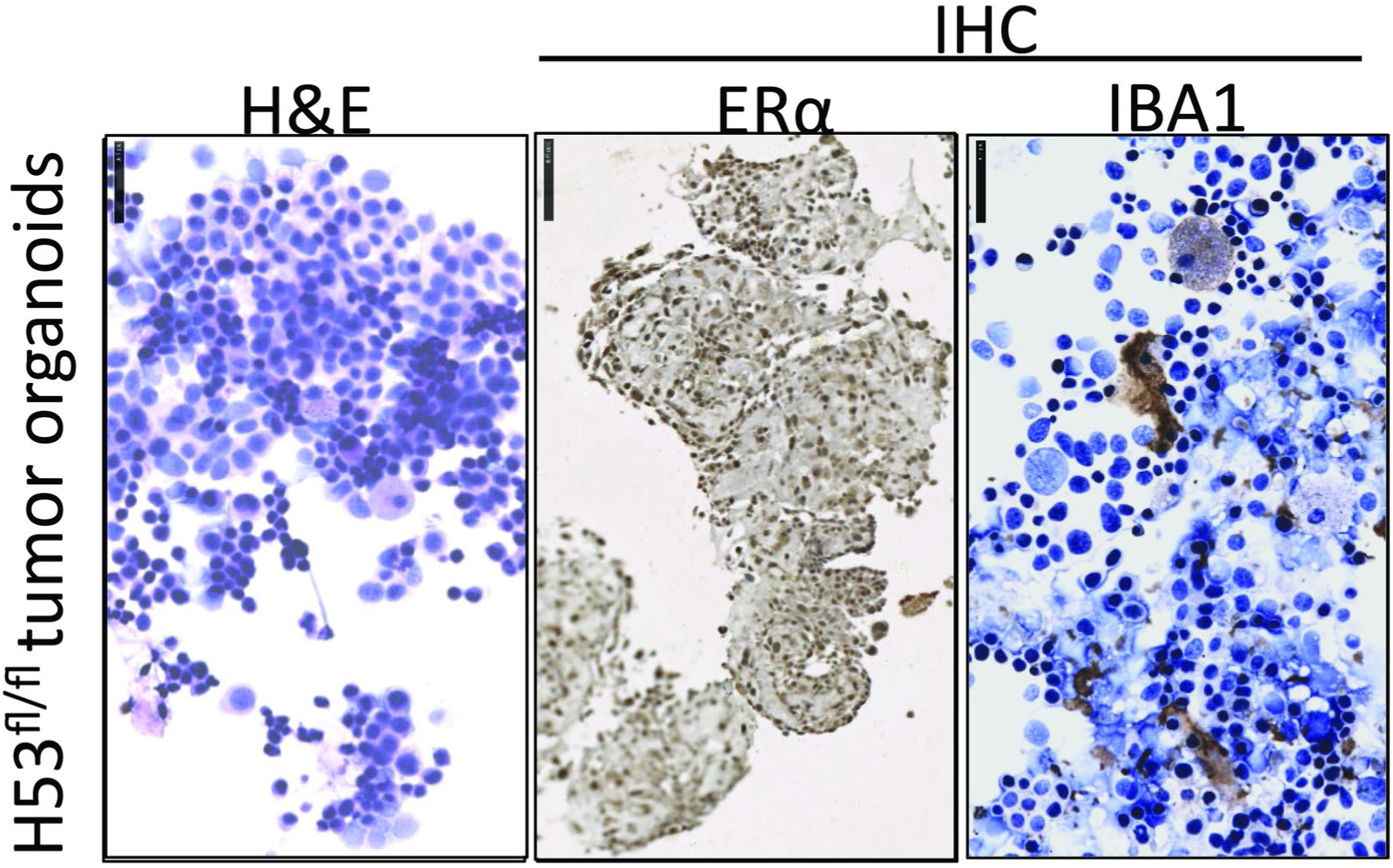
Characterization of murine breast cancer organoids. H&E and IHC stained with ER**α**, IBA1 antibodies in H53fl/fl tumor organoid cells. Scale bars of high-power images are 500 **μ**m.

**Figure S2.**
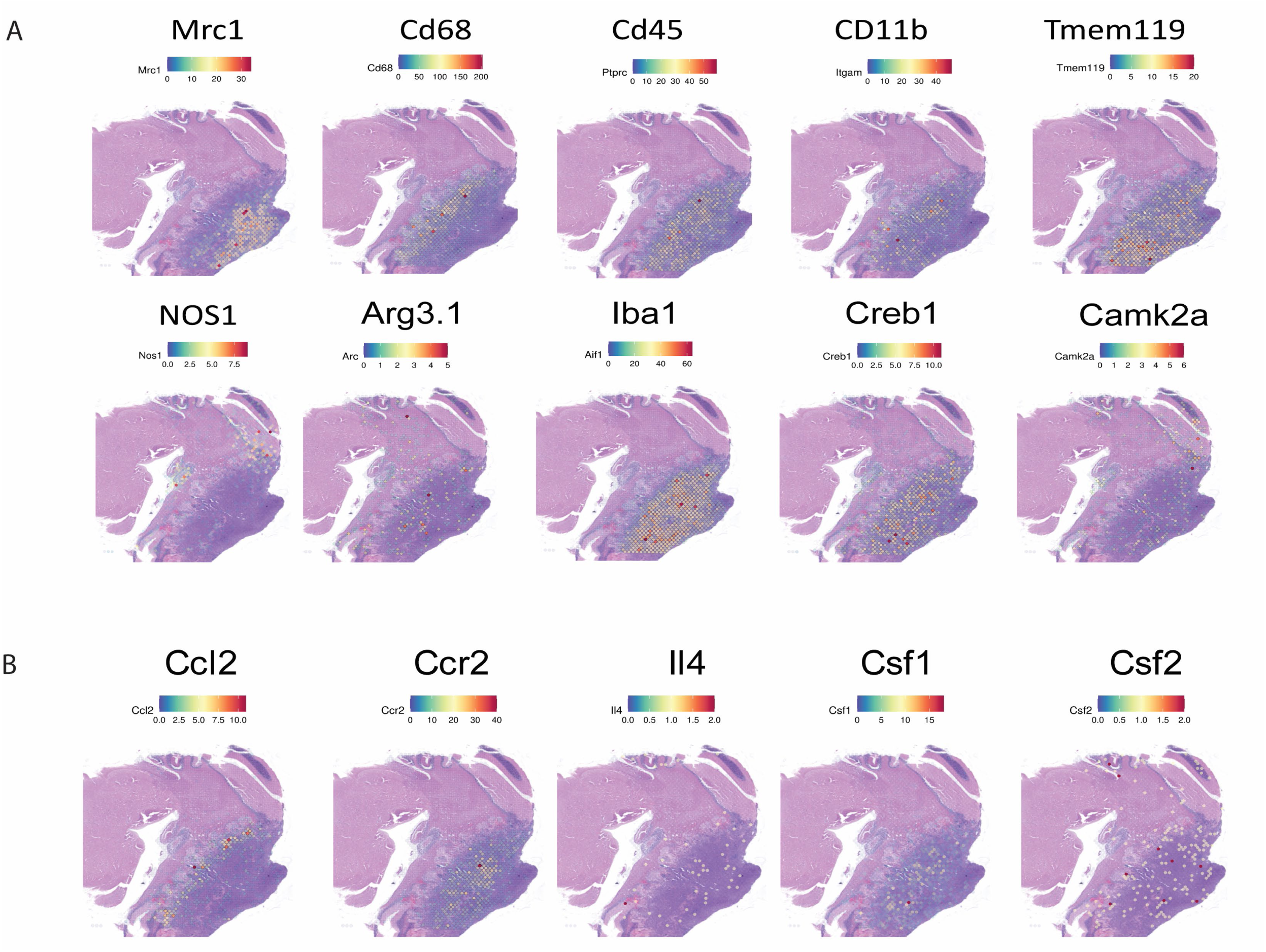
Spatial feature plot of markers for macrophage and its regulation-related chemokines and cytokines. (A) Spatial feature plot of markers for macrophage and microglial cells in the representative tissue sample. (B) Spatial feature plot of key chemokines and cytokines regulate macrophages in the representative tissue sample.

**Figure S3.**
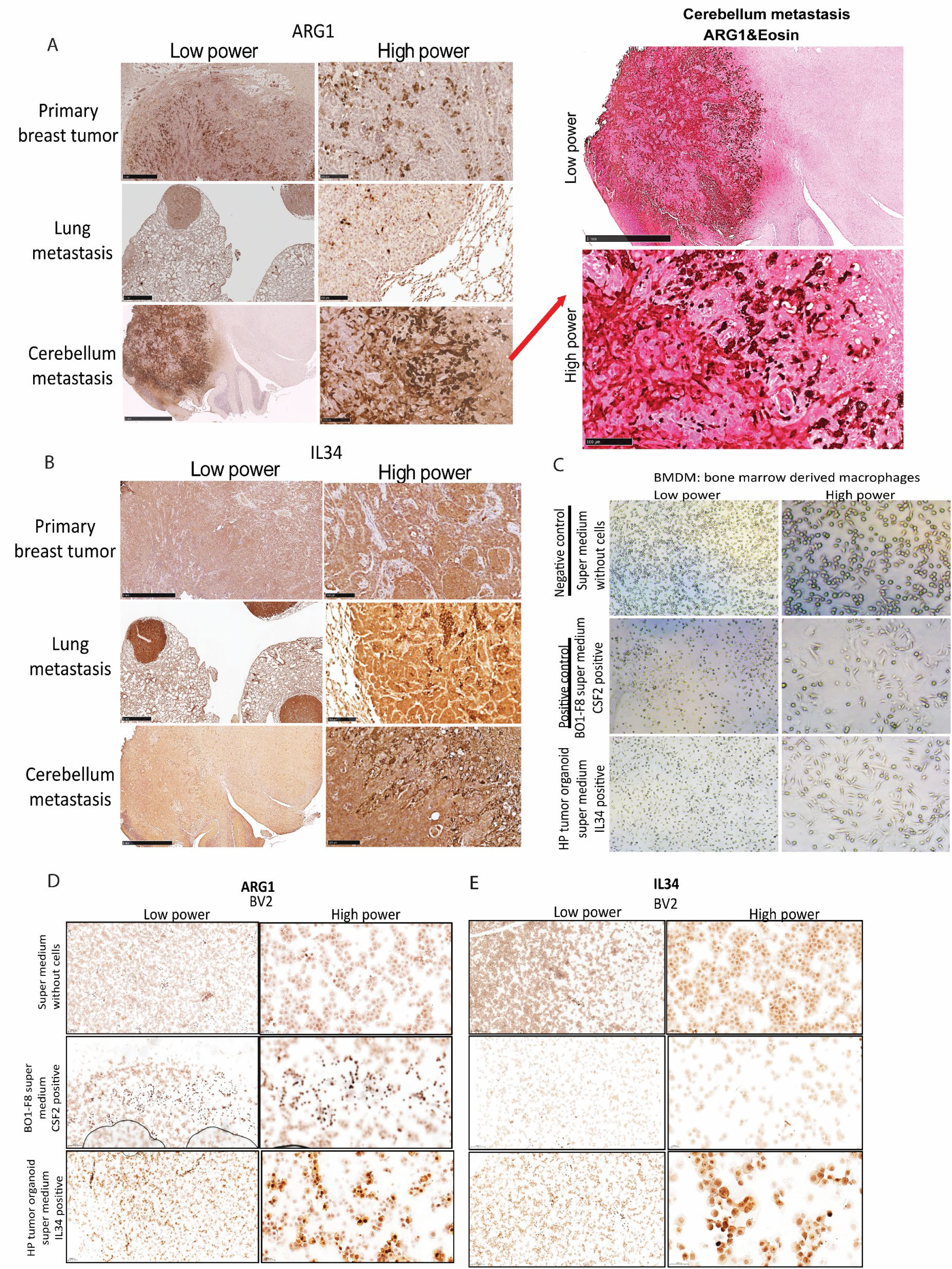
IL34-induced Arg1 expression at the edge of invasion in cerebellum metastasis of HER2+ breast cancer. (A) Left: IHC staining of ARG1 in primary breast tumors, lung metastatic tumors, and cerebellum metastatic tumors from tissue samples. Right: IHC staining of ARG1 and eosin staining in cerebellum metastatic tumors from the tissue sample. (B) IHC staining of IL34 in primary breast tumors, lung metastatic tumors, and cerebellum metastatic tumors from tissue samples. (C) The mouse BMDM was cultured with conditioned media supernatant from BO1^39^ cells or HP tumor organoid cells for 24 hours, and the bright field image shows the cell morphology change. (D) The mouse BV2^40^ was cultured with conditioned media supernatant from BO1^39^ cells or HP tumor organoid cells for 24 hours, and the IHC staining of ARG1 was performed on the cytospin slides of the co-cultured cells. (E) The mouse BV2 was cultured with a conditioned media supernatant from BO1^39^ cells or HP tumor organoid cells for 24 hours, and the IHC staining of IL34 was performed on the cytospin slides of the co-cultured cells.

**Figure S4.**
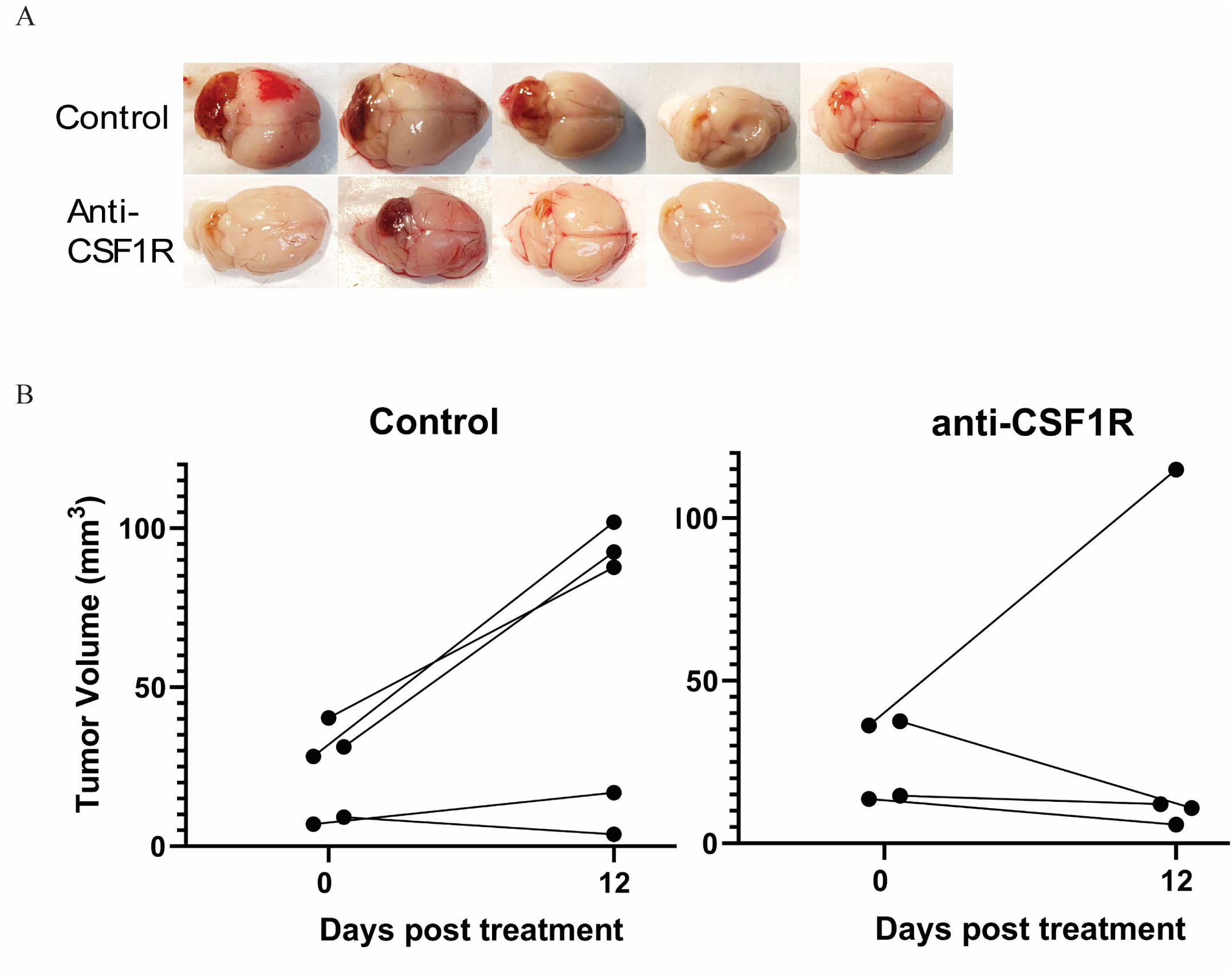
IL34-induced Arg1 expression at the edge of invasion in cerebellum metastasis of HER2+ breast cancer. (A) Brightfield images of dissected cerebellum tumor from control and anti-CSF1R mAb groups. (B) tumor volume change in the control and anti-CSF1R mAb groups measured by MRI at day 0 and day 12 post-treatment.

